# Constraints on the G1/S transition pathway may favor selection of multicellularity as a passenger phenotype

**DOI:** 10.1101/2025.08.20.671217

**Authors:** Tom Louis Ducrocq, Damien Laporte, Bertrand Daignan-Fornier

## Abstract

Multicellularity has emerged in the three branches of the tree of life. The formation of simple multicellular entities can either result from cells aggregating or staying together after mitosis. However, it is not yet fully understood how, once formed, these simple multicellular entities could be maintained or even selected for. Here, using the *ace2* yeast snowflake model of simple multicellularity, we aimed at identifying genetic conditions favoring its maintenance. Growth-competition experiments revealed that, while the *ace2* mutation by itself does not provide any fitness advantage or disadvantage, the *ace2* snowflakes were strongly selected when combined to conditions affecting regulators of the G1/S transition of the cell cycle, such as Cln3 or Whi5. We show that this selection results from a faster exit from quiescence of the *ace2* snowflake cells. Importantly, this advantage is not dependent on the multicellular phenotype, but rather on the *ace2* genotype itself. We found that the *ace2* selective advantage in the *cln3* background fully depends on the *KSS1* gene, a target of the Ace2 transcription factor. Finally, we show that phenotypes observed for *ace2* mutants are phenocopied by the *AMN1^368D^* allelic form found in “non-laboratory” yeast strains, hence adding physiological relevance to these observations. Altogether, our results support the hypothesis that simple multicellularity could, in some cases, persist, not because it provides a direct selective advantage due to multicellularity itself, but rather as a “passenger” phenotype that is maintained alongside other selected traits.

## Introduction

Multicellularity has evolved repeatedly and independently in distinct lineages within bacteria, archaea and eukaryotes (1). This evolutionary process firstly involves the selection of simple forms of multicellularity such as clusters of cells. Several selective drivers of simple multicellularity have been proposed (2) that are generally associated to changes in geometry allowing access to new resources (faster sedimentation, increased mobility, foraging, dispersal), as well as other advantages associated with size increase (predation avoidance, stress resistance, improved extracellular metabolism) and finally, chimerism: the capacity to combine different cellular phenotypes in a single entity, allowing the establishment of intra-organism division of labor.

Many forms of multicellularity are facultative and their formation is triggered by external signals. One of the best studied case is that of the slime mold *Dictyostelium discoideum*, which cells, in response to resource limitation, aggregate to form a fruiting body that favors dispersal of spores (3,4). Another example comes from *Saccharomyces cerevisiae* cells that, under poor nitrogen conditions, can become highly polarized and stay attached, thereby favoring foraging, a phenomenon known as pseudohyphal growth (5,6). In such cases, multicellularity provides alternative options to the non-proliferation observed in unicellular organisms in response to local nutritional starvation (7). These multicellularity-specific responses are triggered by nutritional cues and, some of them may have evolved by functional recruitment of unicellular pre-existing sensors and pathways. It is hence likely that some actors of nutritional control of cell cycle progression should modulate facultative multicellularity. Indeed, there are several experimental evidences connecting the regulators of G1/S transition in the cell cycle to multicellularity. As described below for several species, this mostly concerns two major regulators, retinoblastoma (RB) and Cyclin D. RB is a negative regulator of E2F, a transcription factor critical for G1/S progression, while cyclin D is a negative regulator of RB (8).

Several experimental evidences connect regulators of G1/S transition, and notably RB, to multicellularity. First, when RB from the simple multicellular volvocine *Gonium pectorale* was expressed in its close unicellular relative *Chlamydomonas*, the C*hlamydomonas* cells behaved as a simple multicellular colonial entity (9). This points to an important role for RB in triggering uni- to multicellular transition. Second, in the aggregative multicellular slime mold, *Dictyostelium discoideum*, the RB orthologue, RblA, was found to be involved in the stalk *vs* spore cell-fate preference (10). Finally, another key regulator of G1/S transition, namely Cyclin D, has also been associated to uni- to multicellular transition in two different yeast species, *Candida albicans* and *Saccharomyces cerevisiae* (11,12). Depending on environmental conditions, *Candida albicans* can grow either as a yeast unicellular planktonic form, or as a multicellular hyphal form (13). The transition from one form to the other is associated with invasion and virulence. Strikingly, the knock-down of the *C. albicans* cyclin D orthologue Cln3 resulted in hyphal growth suggesting that the G1 phase of the cell cycle is important in the transition from uni- to multicellular development in *C. albicans* (11). Similarly, in *S. cerevisiae* deletion of *CLN3*, the budding yeast cyclin D homolog, enhanced pseudohyphal growth (12). Interestingly, *CLN3* encodes an inhibitor of Whi5 which is the yeast functional equivalent of RB (13). It hence seems that, during evolution, uni- to multicellularity transitions have repeatedly used pre-existing regulators of the G1/S transition, such as RB and cyclin D. These intriguing observations, from phylogenetically distant organisms, connecting G1/S transition regulators to multicellularity led us to ask the following question. Could the evolution of the G1/S regulation pathway itself have somehow favored multicellularity emergence and/or maintenance?

In this work we used *S. cerevisiae* to question the complex relationship between G1/S cell cycle regulators and multicellularity. We took advantage of the snowflake model (14,15) in which multicellularity is due to a genetic defect in cell separation caused by the deletion of Ace2, a transcription factor that control the expression of chitinases and endoglucanases required for cell wall degradation after cytokinesis completion (16,17). When planktonic *ACE2* and snowflake *ace2* cells were co-cultured without applied selective pressure and their fitness compared, we observed that *CLN3* deletion, as well as *WHI5* overexpression, favored the *ace2* snowflake form. We showed that exit from quiescence is the critical stage leading to the *ace2 cln3* snowflakes selection. Importantly we demonstrated that the *ace2* mutation, but not the snowflake phenotype, was responsible for the *cln3* faster quiescence exit phenotype. Finally, we show that both the multicellularity and quiescence exit phenotypes observed for *ace2* mutants are phenocopied by the *AMN1^368D^*allele found in “non-laboratory” yeast strains. Our results point to the possibility that evolution of cell cycle control could promote the evolution of multicellularity as a passenger phenotype.

## Results

### The *ace2 cln3* snowflake strain is selected over the *ACE2 cln3* planktonic strain

In order to investigate the relationship between the G1/S cell cycle regulators and multicellularity, we used the yeast snowflake model in which the absence of Ace2, a transcription activator of several chitinase and endoglucanases genes (18–20), results in the lack of separation of the daughter cell from the mother cell after cytokinesis. It ensues that *ace2* mutant yeasts form cell clusters that grow and end-up breaking most probably due to mechanical constraints (21). A major advantage of studying snowflakes rather than inducible forms of multicellularity, such as pseudo-hyphae, is that unicellular (planktonic, PK) and multicellular (snowflake, SF) yeast, differing only by one mutation in the *ACE2* gene, can be co-cultivated and their fitness compared. We hence grew isogenic diploid *ACE2*/*ACE2* and *ace2*/*ace2* cells of the BY (S288c-derived) background in minimal enriched medium (SDcasaWAU). Cultures of both genotypes were mixed to obtain a 50/50 ratio of *ace2* SF and *ACE2* PK strains, either at the phenotype level (number of entities observed by microscopy, a SF being an entity with 4 or more cells, Figure 1-figure supplement 1) or at the cell genotype level (by monitoring the *ace2::kanMX* disruption using qPCR) (Figure 1A). The mixtures were then repeatedly grown to saturation and diluted every 2 days, resulting in a total of more than 100 generations. Interestingly, the *ace2* strain showed no selective advantage or disadvantage over the *ACE2* strain, both at the SF phenotype level (Figure 1B, black bars) and the genotype level (Figure 1C, black bars). This indicates that, under these growth conditions, the *ace2* mutation and its multicellular phenotype have no effect on fitness. Strikingly, when similar experiments were performed with *ace2* or *ACE2* strains carrying a deletion of the *CLN3* gene, a deletion known to lengthen the G1 phase and to significantly increase cell volume (22), the *ace2* snowflake strains rapidly took over the population (red bars, Figure 1B and 1C). A similar result was obtained when *WHI5,* encoding the major downstream target of *CLN3*, was overexpressed (red bars, Figure 1D). These results strongly suggest that the observed effect on population fitness takes place through the *CLN3/WHI5* G1/S transition regulatory pathway. Together our results reveal a robust phenotypic interaction between the G1/S transition major players, *CLN3* and *WHI5*, and the *ace2* snowflakes.

**Figure 1.**
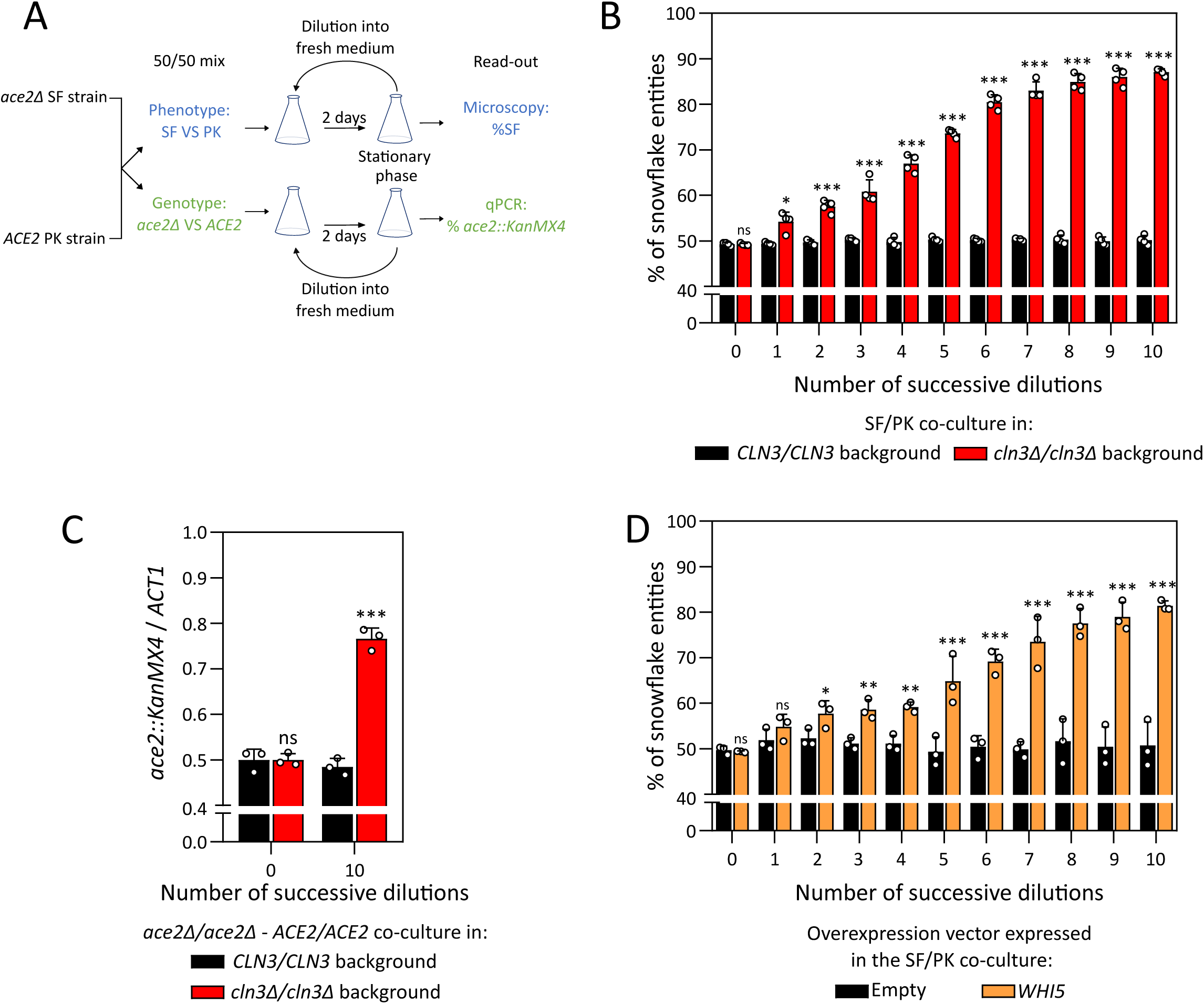
Competition experiments. **(A)** Experimental set-up of the co-culture competition experiments between *ace2/ace2* and *ACE2/ACE2* strains. A 50/50 mix of the two strains based either on phenotype (snowflake vs planktonic, microscopy) or on genotype (*ace2* vs *ACE2,* qPCR) was inoculated in SDcasaWAU liquid medium. The resulting co-culture was grown for 2 days until stationary phase was reached and then diluted in new SDcasaWAU medium. For each dilution step, the proportion of the snowflake phenotype or *ace2* genotype was determined (by microscopy or qPCR respectively). **(B)** Evolution of the percentage of snowflake entities during independent co-culture competitions between *ace2/ace2* and *ACE2/ACE2* strains, either *cln3/cln3* (red bars) or *CLN3/CLN3* (black bars) (N=4, n>250, mean ± SD, ns p>0.05, *p<0.05, ***p<0.001). The percentage of snowflakes is statistically compared between *CLN3/CLN3* and *cln3/cln3* co-cultures at each dilution round, using a Fisher’s exact test. **(C)** The proportion of *ace2/ace2* genotype was monitored by qPCR in independent competitions in *cln3/cln3* (red bars) and *CLN3/CLN3* (black bars) backgrounds. The *ACT1* locus, amplified in all the cells of the population, was used to normalize the proportion of *ace2* cells in the co-culture cell population (N=3, mean ± SD, unpaired t-test, Welch correction, ns p>0.05, ***p<0.001). **(D)** Evolution of the percentage of snowflake entities during independent co-culture competitions between *ace2/ace2* and *ACE2/ACE2* strains overexpressing *WHI5* (orange bars) or not (black bars) (N=3, n>235, mean ± SD, Fisher’s exact test, ns p>0.05, *p<0.05, **p<0.005, ***p<0.001).

### *ace2 cln3* cells exit faster from quiescence than *ACE2 cln3*

In our competition experimental setup (Figure 1A), cultures alternate phases of active cell division with stationary phases during which cells are in a quiescent state (Figure 2-figure supplement 1A). Theoretically, the fitness advantage of the *ace2 cln3* strain over its *ACE2 cln3* isogenic counterpart could be due to a faster doubling time during exponential growth, to a higher biomass yield of the population, to a better survival in stationary phase or to a faster exit from stationary phase, each of these possibilities (schematized Figure 2-figure supplement 1A) acting either alone or in combination. We tested these four possibilities separately. First, we found that the population doubling times of the two sets of strains was not significantly different (Figure 2-figure supplement 1B). In addition, *ace2* and *ACE2* cell populations led to similar biomass accumulation indicating that the yield of carbon utilization was not significantly modified by the *ace2* and/or the *CLN3* mutation (Figure 2-figure supplement 1C). We then estimated cell survival in stationary phase by staining 2 days-old cells with methylene blue, a dye that specifically accumulates in dead cells. No significant differences in dead cell percentages were observed between *ace2* and *ACE2* cells should they be *cln3* or *CLN3* (Figure 2-figure supplement 1D). From these experiments, we conclude that neither the doubling time, nor the biomass yield, or the cell death rate could explain the selective advantage of the *ace2 cln3* cells over their *ACE2 cln3* counterpart.

**Figure 2.**
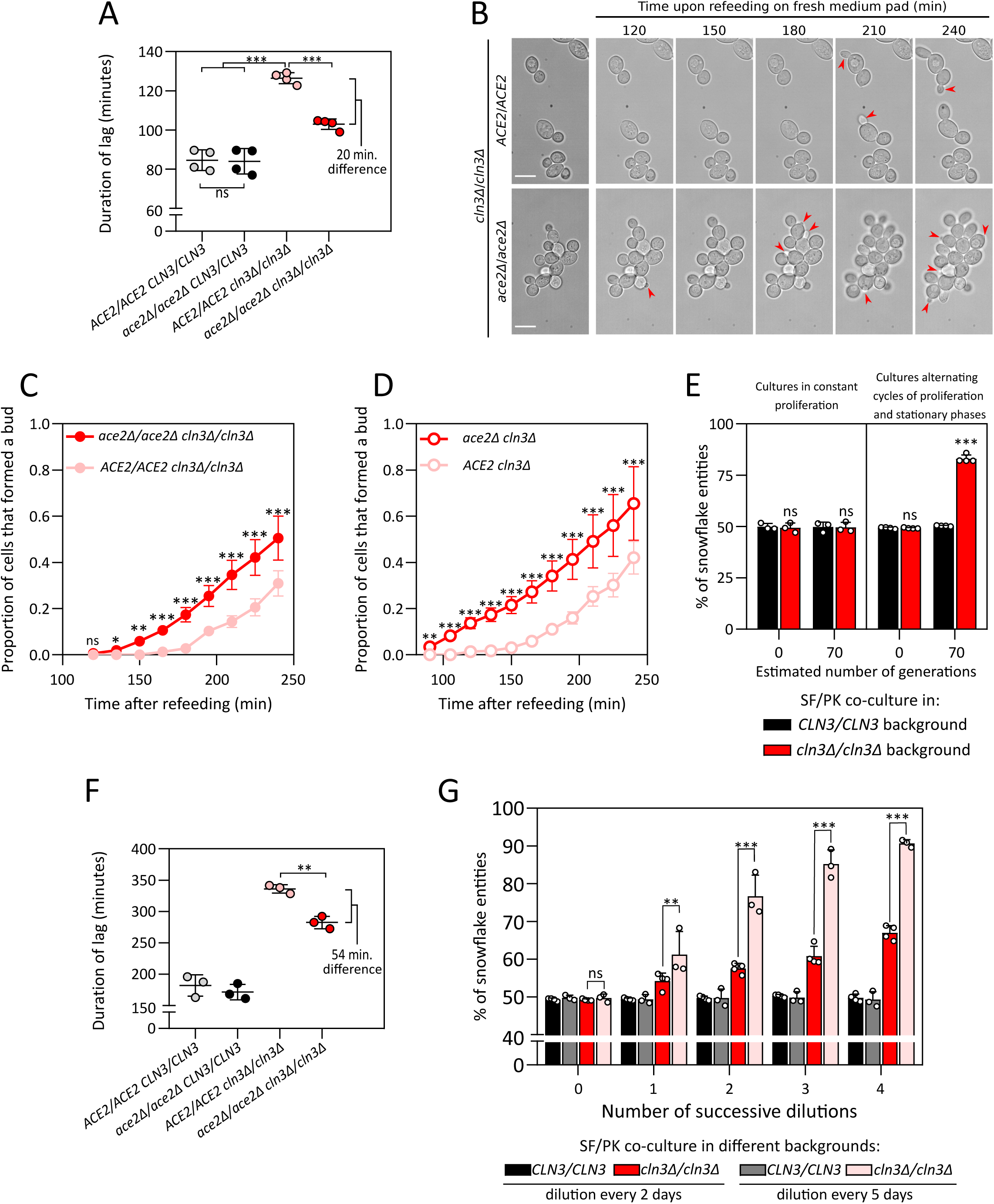
Quiescence exit efficiency explains the results of competition experiments. **(A)** Lag phase duration after cell 2-day old population refeeding with SDcasaWAU medium. The optical density at 600nm was followed after population refeeding for 4 independent experiments. The mean and the SD are indicated together with an unpaired t-test, Welch correction, ns p>0.05, ***p<0.001. **(B)** Representative image series of *cln3/cln3* cells either *ace2/ace2* or *ACE2/ACE2* after refeeding of 2 days cultures on a SDcasaWAU medium containing microscope pad. Arrows point to emerging buds. Bars are 10µm. **(C)** Proportion of *cln3/cln3* cells that have formed a new bud after refeeding on a SDcasaWAU medium containing microscope pad of either *ace2/ace2* or *ACE2/ACE2* 2 days cultures (N=3, n>101, Fisher’s exact test, mean ± SD, ns p>0.05, *p<0.05, **p<0.005, ***p<0.001). **(D)** Same as C in a *cln3* haploid background (N=3, n>100, mean ± SD, Fisher’s exact test, **p<0.005, ***p<0.001). **(E)** Percentage of snowflake entities observed in independent co-cultures of *ace2/ace2* and *ACE2/ACE2* strains. Cultures either went through cycles of proliferation and stationary phase as described in Fig1A (right panel), or maintained in constant exponential phase (left panel) (N>3, n>208, mean ± SD, Fisher’s exact test, ns p>0.05). **(F)** Same as A except that the refeeding was done after a culture of 5 days (N=3, mean ± SD, unpaired t-test, Welch correction, **p<0.005). **(G)** Percentage of snowflake entities observed in independent co-cultures of *ace2/ace2* and *ACE2/ACE2* strains diluted every two days (red and black bars) or every five days (grey and pink bars) (N≥3, n>700, mean ± SD, Fisher’s exact test, ns p>0.05, **p<0.005, ***p<0.001).

We then compared the time required for *ace2 cln3* and *ACE2 cln3* quiescent-cell populations to exit stationary phase. We found that the time required for population to resume proliferation after refeeding, referred to as lag-time, was not significantly different between *ace2 CLN3* and *ACE2 CLN3* cells (grey and black dots, Figure 2A). In addition, we confirmed that the lag-time was significantly longer for the *ACE2 cln3* (light red dots) than for *ACE2 CLN3* cell population (black dots) (Figure 2A), as previously described (23,24). Indeed, after refeeding of stationary phase cultures, *CLN3* transcription is increased by 5- to 10-fold within five minutes (24). Cln3 is hence likely to play an important role for quiescent cells to move out from G1 and reenter proliferation. Importantly, this difference was partially but significantly suppressed in *ace2 cln3* (red dots) cell populations (Figure 2A) suggesting that the *ace2* mutation genetically suppresses the *cln3* strain lag-time defect. We then re-fed quiescent cells on microscope agarose pads containing new medium and addressed cell ability to emit a new bud *i.e.* to exit quiescence. While the time required by cells to exit quiescence was not significantly different between *ace2 CLN3* and *ACE2 CLN3* strains (Figure 2-figure supplement 1E), it was significantly lower for the *ace2 cln3* cells than for the *ACE2 cln3* cells (Figure 2C). Importantly, similar observations were made with haploid *ace2 cln3* cells (Figure 2D) for which snowflakes are much smaller (Figure 2-figure supplement 1G-K). Indeed, not only haploid cells display a smaller cell volume than diploid cells, but their proximal budding pattern make haploid snowflakes much more globular than diploid snowflakes that are rather elongated (25). This observation confirmed the robustness of the *cln3* phenotype and its suppression by *ace2*. It also suggests that this phenotype is not dependent on the size of the snowflake entity. Altogether these results point to quiescence exit as the key step accounting for the competition outcomes.

A simple prediction from these results is that a competition experiment in co-culture ran under conditions where the cells are always maintained in proliferation should not result in any fitness difference between for *ace2 cln3* and *ACE2 cln3* cells. This is what was observed when maintaining these two strains in proliferation for 94hrs (corresponding to at least 70 generations, Figure 2E, left panel), by contrast to an equivalent number of generations obtained by alternating proliferation and stationary phases (Figure 2E, right panel). Reciprocally, as described previously (26), increasing the duration of stationary phase caused an increase of the lag phase length at the population level (Figure 2F). Importantly, it also increased the fitness advantage of *ace2 cln3* over *ACE2 cln3* cells (red and pink bars Figure 2G), while there was still no difference between *ace2 CLN3* and *ACE2 CLN3* cells (black and grey bars Figure 2G). Altogether, our results pinpoint exit from quiescence as the critical stage in which the *ace2 cln3* snowflakes are positively selected.

### The advantage of *ace2 cln3* during quiescence exit is independent of the snowflake phenotype

Why do *ace2 cln3* cells exit quiescence faster than *ACE2 cln3* cells? This could be due to some specificity associated with *cln3* mutation that would modify the snowflake properties. Alternatively, it could be dependent on the specific *ace2 cln3* mutation combination independently of the snowflake phenotype. To answer this question, we dissociated the snowflake phenotype from the *ace2* mutation and vice versa.

First, we asked whether the snowflake phenotype was sufficient, independently of the *ace2* mutation, to rescue the quiescence-exit default associated with *cln3*. Ace2 is a transcription factor that induces the expression of several chitinases and endoglucanases necessary for the septum digestion and cell separation. To mimic the *ace2* snowflake phenotype, we constructed a quintuple chitinase and endoglucanase mutant (*cts1, dse2 dse4, egt2, scw11*), hereafter “quintuple mutant”, that results in a snowflake phenotype comparable to that of the *ace2* mutant (Figure 3-figure supplement 1) while carrying the wild-type *ACE2* allele. When combined with the *cln3* mutation, the five deletions have no effect on quiescence exit efficiency (Figure 3A). Accordingly, the *cln3 ace2* strain exited quiescence much more efficiently than the *cln3* quintuple mutant (Figure 3B). This indicates that the *ace2* mutation itself, and not multicellularity, suppresses the quiescence exit defect of the *cln3* mutant. Finally, competition experiments in co-cultures initially containing equal proportion of quintuple mutant and WT cells confirmed that the snowflake phenotype is not sufficient to give the *cln3* mutant a phenotypical advantage (Figure 3C).

**Figure 3.**
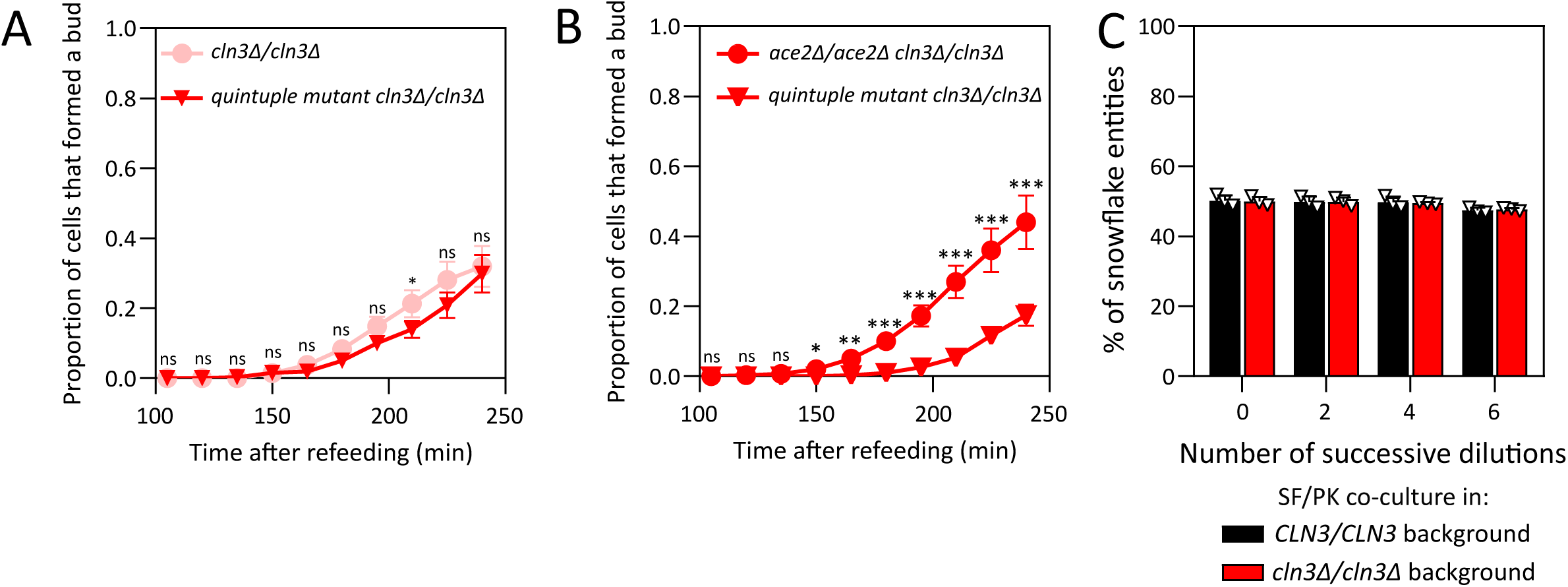
Quintuple *cts1, dse2, dse4, egt2, scw11* mutant strains do not rescue the quiescence-exit default associated with *cln3* and are not selected in competition experiments. **(A)** Proportion of *cln3/cln3* cells, either quintuple mutant or WT, that formed a new bud after refeeding of 2 days cultures on a SDcasaWAU medium containing microscope pad (N=3, n>102, mean ± SD, Fisher’s exact test, ns p>0.05, *p<0.05). **(B)** Proportion of *cln3/cln3* cells, either quintuple mutant or *ace2/ace2*, that formed a new bud after refeeding of 2 days cultures on a SDcasaWAU medium containing microscope pad (N=3, n>75, mean ± SD, Fisher’s exact test, ns p>0.05, *p<0.05, **p<0.005, ***p<0.001). **(C)** Evolution of the percentage of snowflake entities during independent co-culture competitions between quintuple mutant and WT strains either *cln3/cln3* (red bars) or *CLN3/CLN3* (black bars) (N=3, n>250, mean ± SD, Fisher’s exact test).

Reciprocally, we asked whether the *ace2* mutation itself, independently of the snowflake phenotype, could be sufficient to rescue the quiescence-exit default associated with *cln3*. We thus examined whether planktonic cells obtained in an *ace2* mutant background would still behave as their snowflake counterparts. To obtain planktonic *ace2* cells, we used sonication to physically break the snowflakes, before triggering quiescence exit. Mild sonication resulted in an increased proportion of planktonic cells in the *ace2 cln3* cell population (up to 50%, Figure 4-figure supplement 1A) with no defect in cell survival (Figure 4-figure supplement 1B). After sonication, isolated cells, small and large cell clusters displayed a similar quiescence exit efficiency (Figure 4A). Furthermore, the sonicated and non-sonicated cells had the same quiescence exit efficiency, indicating that sonication *per se* had no significant impact (Figure 4-figure supplement 1C). Still, after sonication, *cln3 ace2* isolated cells exited quiescence faster than their *cln3 ACE2* counterpart (Figure 4B). These results show that the snowflake phenotype is not necessary for the ability of the *ace2 cln3* mutant to exit quiescence faster than *ACE2 cln3* cells.

**Figure 4.**
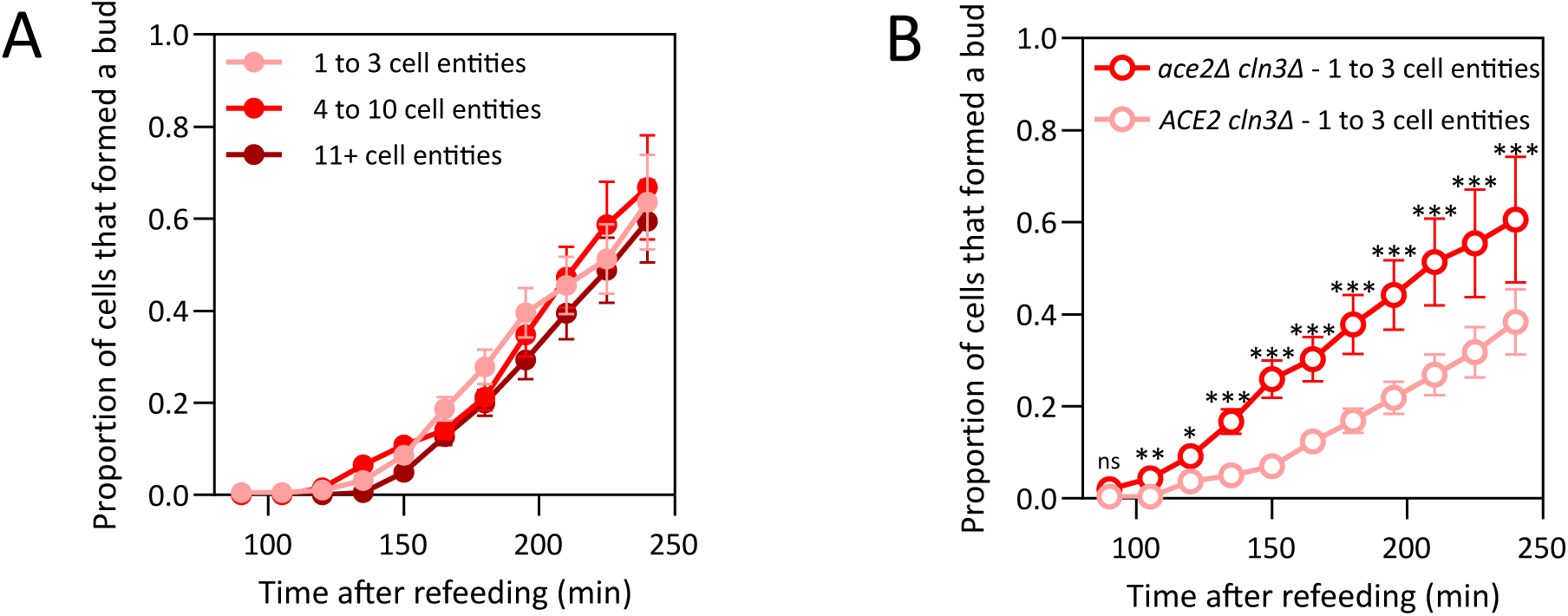
Rescue of the quiescence-exit default associated with *cln3* depends on the *ace2* mutation and not on the Snowflake phenotype. **(A)** A 2-day *ace2/ace2 cln3/cln3* culture was sonicated and refed onto a fresh SDcasaWAU microscope pad. The proportion of cells that formed a new bud after refeeding on a SDcasaWAU medium containing microscope pad was assessed within different multicellular size ranges (N=3, n>184, mean ± SD, Fisher’s exact test). **(B)** 2-day *cln3* cultures, either *ace2* or *ACE2*, were sonicated and refed onto a fresh SDcasaWAU microscope pad. Within entities formed of 1 to 3 cells, the proportion of cells that formed a new bud after refeeding on a SDcasaWAU medium containing microscope pad was assessed (N=3, n>63, mean ± SD, Fisher’s exact test, ns p>0.05, *p<0.05, **p<0.005, ***p<0.001).

Together, our results with the haploid strains, the quintuple mutant and sonication experiments establish that the *ace2* mutation, but not the snowflake phenotype, is responsible for the *cln3* fast quiescence exit phenotype.

### The *ace2 cln3* fast quiescence exit phenotype is dependent on the Kss1 MAP kinase

As previously mentioned, a delay of *cln3* mutants to exit quiescence was already documented (24). Why is this phenotype suppressed by the *ace2* deletion? Among Ace2 well-established targets, *AMN1*, *BUD9, CHS7, CTS1, DSE2, DSE4* and *SCW11* are activated, while *KSS1* is down regulated by Ace2 (18). Four of those, namely *CTS1, DSE2, DSE4* and *SCW11*, are not involved in suppressing the delay phenotype since they are mutated in the quintuple mutant that, in combination with *cln3*, has the same delay as a single *cln3* mutant (Figure 3D). We focused on Kss1, a MAP kinase involved in filamentous growth and mating (27,28), because the Kss1 pathway was previously found to positively regulate *CLN1,* a G1 cyclin encoding gene that is partially redundant to *CLN3* (29). We hypothesize that deletion of *ACE2* would result in higher *KSS1* expression (18) that in turn, could lead to higher *CLN1* expression and thereby be responsible for the *cln3* mutation suppression. We hence tested whether suppression of *cln3* by *ace2* was dependent on *KSS1* by comparing quiescence exit efficiency of *ace2 cln3 KSS1* and *ace2 cln3 kss1* cells. Clearly, the knock-out of *KSS1* totally abolished the phenotypic suppression of *cln3* by *ace2* (Figure 5A). Further, *CLN1* expression upon quiescence exit was significantly higher in the *ace2 cln3* than in the *ACE2 cln3* strain (Figure 5B), this difference of expression being entirely abolished in the absence of *KSS1* (Figure 5C). We conclude that the competitive advantage of the *ace2* mutant over its *ACE2* wild-type counterpart is most likely due to higher expression of *CLN1* that, at least partially, compensates for the absence of *CLN3*. Importantly, this transcriptional effect of Ace2 on *CLN1*, *via* Kss1, is independent of the role of Ace2 on multicellularity.

**Figure 5.**
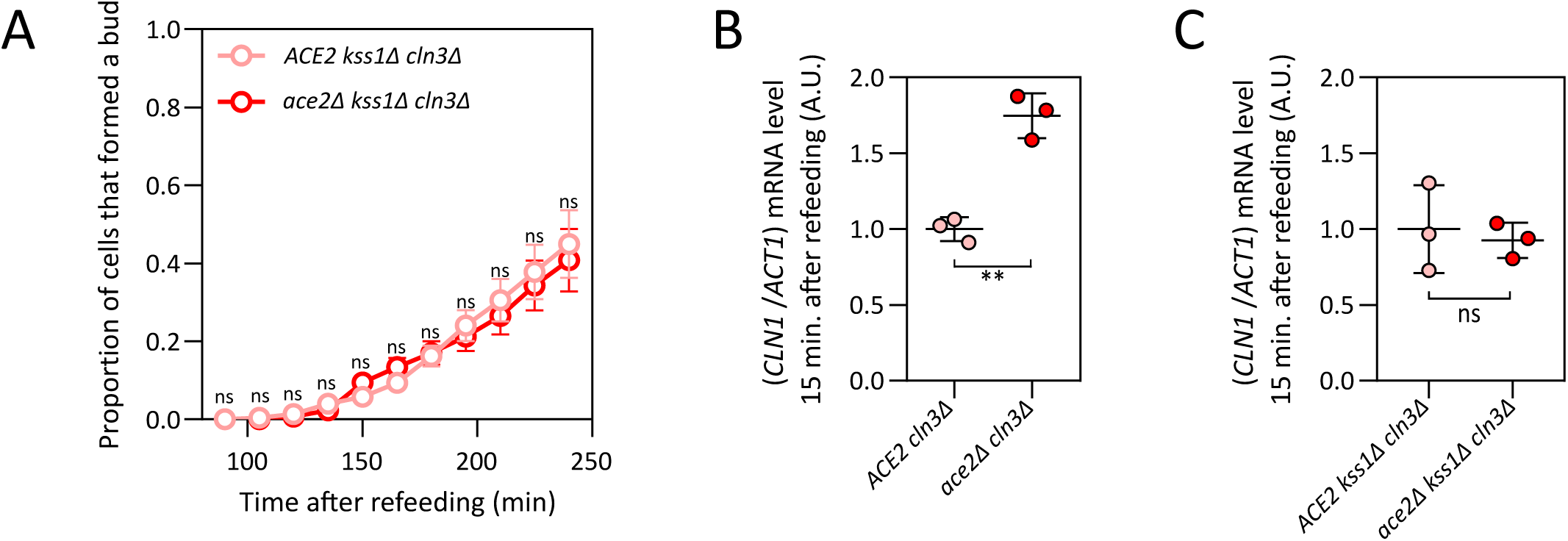
Rescue of the quiescence-exit default associated with *cln3* is dependent on the Kss1 MAP kinase, which favors transcription of the *CLN1* cyclin gene. **(A)** Proportion of *cln3 kss1* cells, either *ace2* or *ACE2*, that formed a new bud after refeeding of 2-day cultures on a fresh SDcasaWAU medium containing microscope pad (N=3, n>86, mean ± SD, Fisher’s exact test, ns p>0.05). **(B)** Ratio of *CLN1* mRNA level and *ACT1* mRNA level, determined by RT-qPCR, 15 minutes after refeeding of 2 days *cln3* cultures, either *ace2* or *ACE2* (N=3, mean ± SD, unpaired t-test, Welch correction, **p<0.005). **(C)** Ratio of *CLN1* mRNA level and *ACT1* mRNA level, determined by RT-qPCR, 15 minutes after refeeding of 2-day *kss1 cln3* cultures, either *ace2* or *ACE2* (N=3, mean ± SD, unpaired t-test, Welch correction, ns p>0.05).

### The *AMN1^368D^* allelic-form found in the wild, phenocopies the *ace2* deletion

In laboratory strains, the Ace2 protein level is constant throughout the cell cycle, while it is not always the case in non-laboratory strains (25). This is due to the presence of the recessive *amn1^368V^* allelic form in laboratory strains (hereafter *amn1-368V*), which was probably initially selected to avoid “unwanted” cell clusters. In non-laboratory strains, the Amn1-368D protein induces the proteolysis of Ace2 through the ubiquitin proteasome pathway (25). It thus appears that the Amn1/Ace2 tandem has an important physiological role associated with optional post-mitotic cell separation. We first observed that expression of *AMN1^368D^*in the BY background resulted in a snowflake phenotype comparable to that of *ace2* mutant strains (Figure 6-figure supplement 1). We then asked whether *AMN1^368D^*allelic form could suppress the quiescence exit delay of the *cln3* mutant. Indeed, the *cln3 AMN1^368D^* strain exited quiescence much more rapidly than the *cln3 amn1-368V* strain (Figure 6A). Finally, we found that the *AMN1^368D^* strain was strongly enriched in competition experiments when associated with a *cln3* mutation (Figure 6B) or with *WHI5* overexpression (Figure 6C). We conclude that, in haploid yeast, the *AMN1^368D^* allelic-form phenocopies both the multicellularity and the suppression phenotypes of the *ace2* mutant.

**Figure 6.**
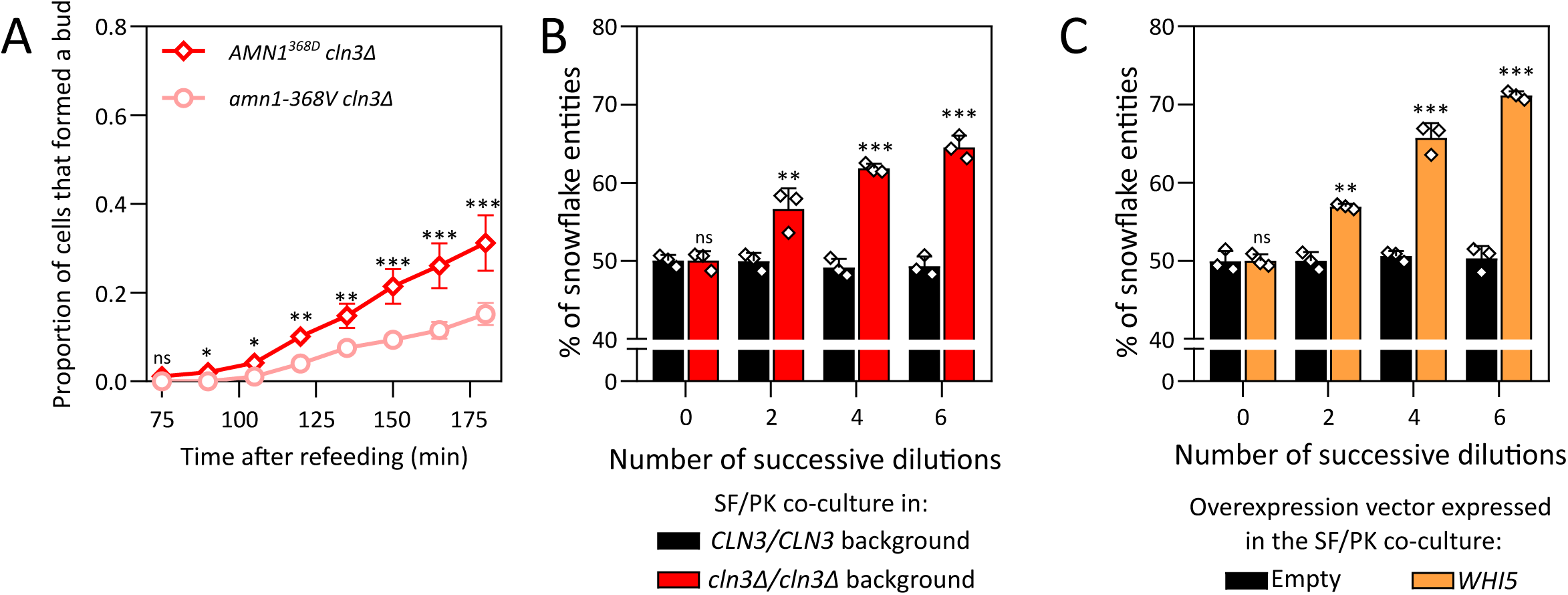
Quiescence exit efficiency and selection in competition experiments of Snowflake *AMN1^368D^* strains. **(A)** Proportion of *cln3* cells, either *AMN1^368D^* or *amn1-368V*, that formed a new bud after refeeding of 2 days cultures on a fresh SDcasaWAU medium containing microscope pad (N=3, n>77, mean ± SD, Fisher’s exact test, ns p>0.05, *p<0.05, **p<0.005, ***p<0.001). **(B)** Evolution of the percentage of snowflake entities during independent co-culture competitions between *AMN1^368D^* and *amn1-368V* strains, either *cln3/cln3* (red bars) or *CLN3/CLN3* (black bars) (N=3, n>252, mean ± SD, Fisher’s exact test, ns p>0.05, **p<0.005, ***p<0.001). **(C)** Evolution of the percentage of snowflake entities during independent co-culture competitions between *AMN1^368D^*and *amn1-368V* strains overexpressing *WHI5* (orange bars) or not (black bars) (N=3, n>302, Fisher’s exact test, ns p>0.05, **p<0.005, ***p<0.001).

## Discussion

In this work, we took advantage of the snowflake model of multicellularity that is genetically determined, by *ace2* or *AMN1* and is hence not triggered by external cues, as it is often the case for facultative multicellularity. This experimental design allows direct fitness measurements between uni-and multicellular entities. Importantly, in a wild-type background, multicellularity was neither selected nor counter-selected, even following a substantial period of co-culture. This establishes that simple multicellular cluster formation should not always come at a cost, contrary to the general assumption in the literature (30,31). Such a neutral effect of multicellularity on fitness could easily result in phenotypical *aller-retours* between unicellularity and multicellularity during evolution. In fact, such *aller-retours* are observed during the life cycle of non-laboratory strains in which diploids are planktonic while haploid strains form clusters (32). Here we show that the evolution of the cell cycle regulation may interact with the multicellular phenotype. We revealed a genetic connection between major G1/S regulators and *AMN1* and *ACE2*, key actors modulating cell cohesion at the end of mitosis (Figure 7). Interestingly, this connection operates differently in laboratory strains carrying the *amn1-368V* allele (Figure 7A) and strains carrying the *AMN1^368D^* allele (Figure 7B). We established that this genetic interaction is due to the pleiotropy of the *ace2* mutation, affecting not only the chitinases and endoglucanases that are critical for septum digestion and cell separation, but also the Kss1 MAP kinase which contributes to the activation of Cln1, one of the G1 cyclins. This phenotypical link between *CLN1* expression and multicellularity, achieved by Ace2 function, is particularly important during quiescence exit, a situation that is likely to happen very often under ecological conditions, such as boom and bust cycles when moving from rotting fruit to rotting fruit. We propose that in some instances, multicellularity may have evolved as a side effect, named here “passenger phenotype”, of another phenotype that was the one selected (Figure 7B). Since transcription factors often have multiple targets, an alteration in their expression or function leading to the selection of a phenotype due to one of the targets could easily lead to the selection of additional phenotypes due to other targets. This could be a common feature involved in evolutionary processes.

**Figure 7.**
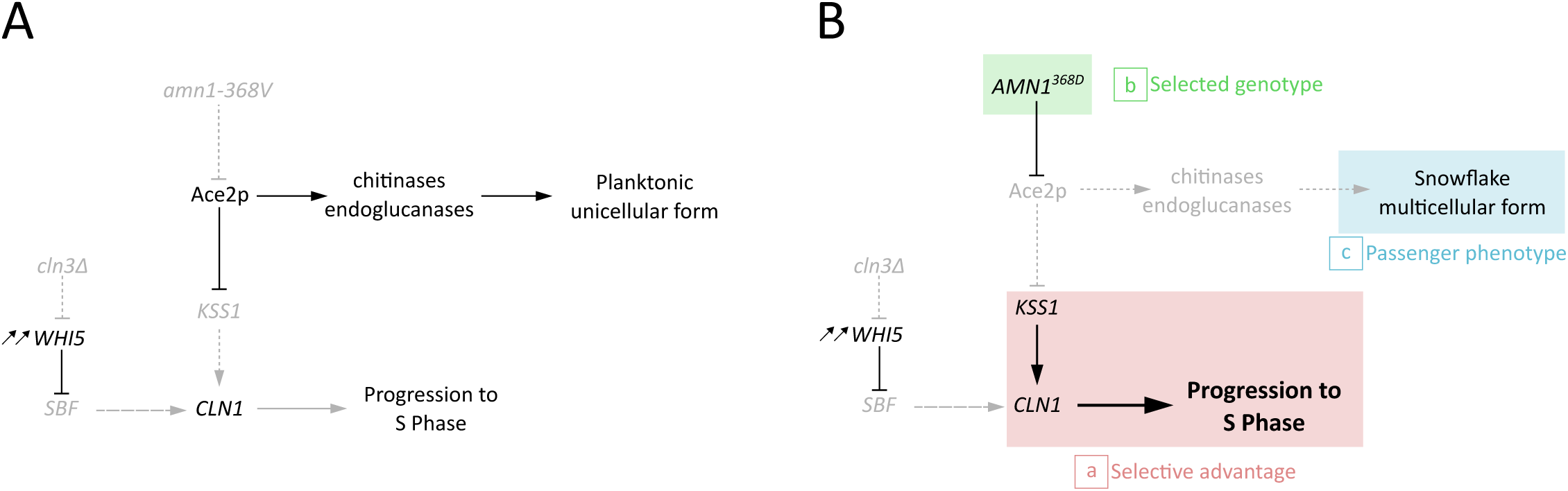
Model explaining the selection of the Snowflake phenotype as a passenger phenotype. In strains expressing the non- functional *amn1-368V* allele **(A)**, a default in quiescence exit after refeeding was observed in a *cln3* mutant. In non-laboratory strains **(B)**, the functional *AMN1^368D^* allele leads to degradation of Ace2p, which can no longer repress the expression of the Kss1 Map kinase. Kss1p favors the transcription of *CLN1*, which would favor progression to S phase and partially suppress the default in quiescence exit associated with the *cln3* mutation. This observed selective advantage (a) leads to the selection of *AMN1^368D^* genotype (b), and because of the pleiotropy of Ace2p transcription factor, to the selection of the Snowflake multicellular phenotype as a passenger phenotype (c).

Connections between G1/S regulators and multicellularity have been repeatedly observed in phylogenetically distant species (see introduction section), suggesting a propitious genetic interplay for co-evolution. Olson and co-workers have shown that RB could have played a role in early steps of multicellularity evolution *via* a cooption mechanism favoring the appearance of simple multicellular entities (9). Our work further supports a role for G1/S regulators in evolution of multicellularity, but the mechanism appears different and affects selection of multicellularity rather than its initial appearance. We show that the *ace2* or *AMN1^368D^* allelic forms are sufficient to counteract a negative fitness effect of the overexpression of the RB functional orthologue Whi5 (Figure 1D, Figure 6F). Importantly we have shown that this phenotypic suppression is independent of the multicellular phenotype associated with the *ace2* or *AMN1^368D^* allelic forms. This observation led us to propose that some regulatory effects on the G1/S transition could be alleviated by Amn1 degradation of Ace2 and thereby connected to multicellularity. In this hypothesis the selective pressure applies on G1/S transition efficiency and not on multicellularity itself. It is hence different from the cooption hypothesis in which the RB protein acquired new functionalities leading to multicellularity that would result in some directly selectable fitness advantage. Both scenarios imply co-evolution of cell cycle regulation and multicellularity, which could therefore be a much more general feature than first thought. It could indeed be a case of convergent evolution as suggested by the fact that it may involve distinct mechanisms.

More generally, our results prompted us to reconsider whether simple multicellularity, which appeared multiple times in evolution, was always selected for the new physical and physiological properties it provides (2). Alternatively, it is possible that multicellularity was initially selected as a byproduct, allowing it to be maintained although not necessarily increasing fitness as such. This would contribute to disconnect the initial selection of simple multicellularity from the selection of further associated advantages, such as division of labor, that would require time and necessitate that the multicellularity phenotype would not be counter-selected in the first place. Such a passenger selection of multicellularity would make it independent of physical constraints such as size or shape that could restrain its further evolution. The work presented here provides a proof of concept that such a scenario is plausible and may have contributed to the multiple appearance of multicellularity in the tree of life.

## Materials and methods

### Yeast media and strains, plasmids and oligonucleotides

Yeast cells were grown in liquid or solid media at 30 °C. SD is a synthetic minimal medium containing 0.5% ammonium sulfate, 0.67% yeast nitrogen base (Difco), 2% glucose. SDcasaWAU is an SD medium supplemented with 0.2% casamino acids (Difco) tryptophan (0.2 mM), adenine (0.3 mM) and uracil (0.3 mM). Unless specified, cells were grown in SDcasaWAU medium. When indicated, medium was depleted in uracil, resulting in a medium named SDcasaWA, allowing the selection of cells expressing a vector carrying *URA3* marker. SC medium was prepared as described by Sherman (33) and was supplemented by histidine (0.06 mM), leucine (0.5 mM), lysine (0.07 mM), tryptophan (0.2 mM), adenine (0.3 mM), and uracil (0.3 mM). The medium obtained by the depletion of both uracil and leucine, called SC-U-L medium, allows the selection of cells expressing both *URA3* and *LEU2* markers. The YPD medium contained 1% yeast extract, 2% peptone, and 2% glucose.

Yeast strains (listed in Table 1) are derived from both FY4 and FY5 prototrophic strains and from disrupted strains isogenic to BY4741 or BY4742 purchased from Euroscarf (Frankfurt, Germany). Strains described in Table 1 were either obtained by crossing, sporulation, and micromanipulation of meiosis progeny or by transformation of cells with a DNA fragment of interest (plasmid, PCR product). Yeast strains were transformed using the lithium acetate method (34).

All plasmids and oligonucleotides are listed in Table 2 and Table 3, respectively. To construct the plasmids allowing the overexpression of *WHI5* in yeast strains, the open reading frame of the *WHI5* gene, driven by its own promoter, was amplified using the genomic DNA of the FY4 strain as a template. The PCR product was then cloned into two 2µ vectors, YEpLac195 and YEpLac181, carrying respectively *URA3* and *LEU2* markers. Two plasmids carrying respectively two different alleles of the *AMN1* gene were also constructed. The open reading frame of the *AMN1* gene was amplified using either the FY4 strain genomic DNA (*amn1^368V^* PCR product) or the GN-1C strain genomic DNA (*AMN1^368D^* PCR product). The GN-1C strain was a generous gift from Pr. Marullo. The PCR products were cloned into YIpLac211, that allowed integration of the cloned *AMN1* allele at the yeast *AMN1* locus.

### Description of a Snowflake population

Snowflake populations or co-culture between Snowflake and Planktonic strains were observed in a Kova Glasstic slide (Kova international, 87144F). The proportion of the Snowflake phenotype in these populations was determined by counting entities formed by four or more cells. To characterize the size of Snowflakes within a population, the population was observed between slide and coverslip separated by a layer of double-sided adhesive tape. The longest segment of the Snowflake entity was measured, as well as the longest segment perpendicular to the first segment and lying in the same Z plane.

### Competitions experiment in co-culture

Snowflake strains (either *ace2* or quintuple mutant or *AMN1^368D^*) and planktonic strains were grown in monocultures to stationary phase, for 2 days unless specified. Snowflake and planktonic strains were then mixed in indicated proportion and co-cultured. Unless specified, every 2 days, co-cultures, after they reached stationary phase, were diluted by a factor of 1000 into fresh liquid medium (10μl in 10ml).

To set up the co-cultures for which the proportion of Snowflake phenotype was monitored, the entity (Snowflake entity or single-cell entity) concentrations in each of the monocultures were determined. The volumes of the 2 monocultures in the mix were accordingly determined and then adjusted to obtain 50% of Snowflake entity in the mix. The proportion of Snowflakes in the co-culture was determined every two days before dilution in fresh medium.

To set up the co-cultures for which the proportion of *ace2* genotype was monitored, the two strains were mixed to obtain a 50/50 cell ratio. Every 2 days, the genomic DNA of the stationary phase co-cultures was extracted by phenol-chloroform protocol. Quantitative PCR amplifications were then performed with GoTaq® qPCR Master Mix (Promega, A6001), using 50ng of DNA extracted from co-cultures. The first set of primers was designed to amplify the *ace2::KanMX4* locus with the first primer targeting the region directly upstream of the ATG of the *ACE2* gene, and the second primer targeting a region specific to the *KanMX4* marker (Supplemental Table 3). The second set of primers targeted a region of *ACT1* gene (Supplemental Table 3).

For competitions in which the co-cultures were maintained in proliferation phase for 94 hours, the co-cultures were diluted every 8 hours to ensure that the OD_600nm_ was always maintained below 0.5.

### Population measurement

To determine the population doubling time of yeast strains, cells were kept in exponential proliferation phase (OD_600nm_≤1.0) for at least 24 hours by successive dilutions before any measurement. Population doubling time was then determined by following the optical density at 600 nm during 5 hours on at least three independent exponential growth culture. Population doubling time was then determined by exponential growth regression of the proliferation curves using GraphPad Prism 8 (GraphPad Software, Inc. La Jolla, USA).

To determine cell viability, the cells were stained with a solution containing 0.2% of methylene blue (Sigma-Aldrich, Saint Louis, MI, USA) and 2% sodium citrate pH7. After 5 minutes of incubation, the proportion of blue-stained dead cells was scored manually.

The biomass yield of cultures was determined by filtration of 2-day stationary phase cultures on 0.45 µm polyamide pre-weighted filters (25006–47-N; Sartorius). The filter was washed twice with 10 mL of miliQ water and then dried with successive 30 seconds cycles in a microwave oven at 800 W until the mass of the filter and cells was constant.

Cell volume was measured by microscopy using the spheroid formula: 4/3*π*a^2^*c, where a and c are the equatorial and polar radius respectively.

### Refeeding of stationary phase cultures

Unless the age of cultures is otherwise specified, 2-day stationary phase liquid monocultures were refed either on liquid or solid fresh SDcasaWAU medium.

Stationary phase cultures were refed by dilution into liquid fresh SDcasaWAU medium at approximately 0.05 OD_600nm_. The OD_600nm_ was monitored upon refeeding for at least 7 hours covering both ‘lag’ and exponential phases (until at least a 5-fold increase of the OD_600nm_ compared to the initial point). To quantify the duration of the ‘lag’ phase, we normalized all the OD_600nm_ values of each individual growth curve to their respective initial OD_600nm_ value and then applied the logarithmic function to all the normalized OD_600nm_ values of the growth curve. We next performed a linear regression on the points corresponding to the proliferation phase of the growth curve using GraphPad Prism 8 (GraphPad Software, Inc. La Jolla, USA). The time value for which the linear regression intersects the Time-X axis corresponds to the end of the ‘lag’ phase and the beginning of the proliferation phase, and is referred as ‘the duration of lag phase’.

Quiescent cells from stationary phase cultures were also refed onto solid medium. Cells were spread onto half a 3% agarose microscope pad made of fresh SDcasaWAU. Bright field images of individual cells were then captured every 15 minutes for up to 4 hours. The cells forming an unequivocal new bud upon refeeding were scored.

When indicated, an aliquot of stationary phase culture can be sonicated before a refeeding experiment on microscope agarose pad. Cycles of 30 seconds in a sonication bath were carried out until the proportion of single-cell entities in the *ace2* SF culture increased to a desired proportion (approximately 40%).

### Determination of *CLN1* mRNA level upon refeeding by Reverse transcriptase-PCR

To determine the level of *CLN1* mRNA upon refeeding of 2-day stationary phase cultures, 15 mL of a 2-day culture was diluted in 85 mL of fresh medium. Fifteen minutes upon refeeding, 25 mL of culture was sampled and centrifuged at 4000 rpm at 4°C for 3 min. The pellet was washed in 1 mL of RNase-free H2O, and centrifuged at 12,000 g at 4°C for 30 seconds. A Tri-Reagent (Sigma, T9424) protocol was then performed to extract total RNA. A reverse-transcription of 200ng of total RNA using the ‘High Capacity cDNA Reverse Transcription’ Kit (ThermoFisher, 4368813) was then performed. The cDNA was diluted to 1/25 and amplified with a Quantitative-PCR protocol, using the GoTaq® qPCR Master Mix (Promega, A6001) and primers specific to the coding region of *CLN1* or the coding region of *ACT1* (Supplemental Table 3).

### Microscopy

The refeeding experiments on microscope pad, as well as the Snowflake size-measurement experiments were carried out using a Zeiss 200M inverted microscope (Carl Zeiss, Thornwood, NY, USA) with the additional equipment : a MS-2000 stage (Applied Scientific Instrumentation, Eugene, OR, USA), a Lambda LS 300 W xenon light source (Sutter, Novato, CA, USA), a 5 positions filter turret, a 100X 1.4NA Plan-Apochromat objective and a CoolSnap HQ camera (Roper Scientific, Tucson, AZ, USA). The filters are from Chroma Technology Corp. The microscope and additional equipment were controlled by SlideBook software 5.0. (Intelligent Imaging Innovations, Denver, CO, USA). The images were analyzed with Image J. All other microscopy experiments were carried out using Zeiss Primo Star microscope, equipped with a 40X/0.65 Plan-Achromat objective.

### Statistical analysis

All experiments were carried out at least three times on independent biological samples. The exact number of independent samples is indicated in the figure legends. Data are presented as mean ± SD. All the statistical analyses were done using GraphPad Prism 8 (GraphPad Software, Inc. La Jolla, USA). The proportion of SF in competition experiments, the refeeding experiments on microscope pad and the proportion of dead cells were analyzed by Fisher’s exact test on a contingency table. Other experiments are analyzed by Welch’s unpaired t-test. p-values below 0.05 are represented by *; p values inferior to 0.005 are represented by **; and p values inferior to 0.001 are represented by ***. p-values superior to 0.05 were considered non-significant and are represented by ‘ns’.

## Acknowledgments and funding sources

The authors thank Benoît Pinson for technical advising, the members of the Sagot lab for discussions on the project and Isabelle Sagot for critical reading of the manuscript. The authors also thank Sabine Vaur for her help with the qPCR method. This work was supported by a grant from Ligue Régionale Contre le Cancer (2022 Dordogne) to BD-F. TD was supported by a grant from University of Bordeaux, NewMoon RRI Program on Cancer in Evolution.

## Author contributions

T.L.D., D.L. and B.D.-F. designed research; T.L.D. performed research; T.L.D., D.L. and B.D.-F. analyzed data; and T.L.D. and B.D.-F. wrote the paper.

---

**Figure 1-figure supplement 1.**
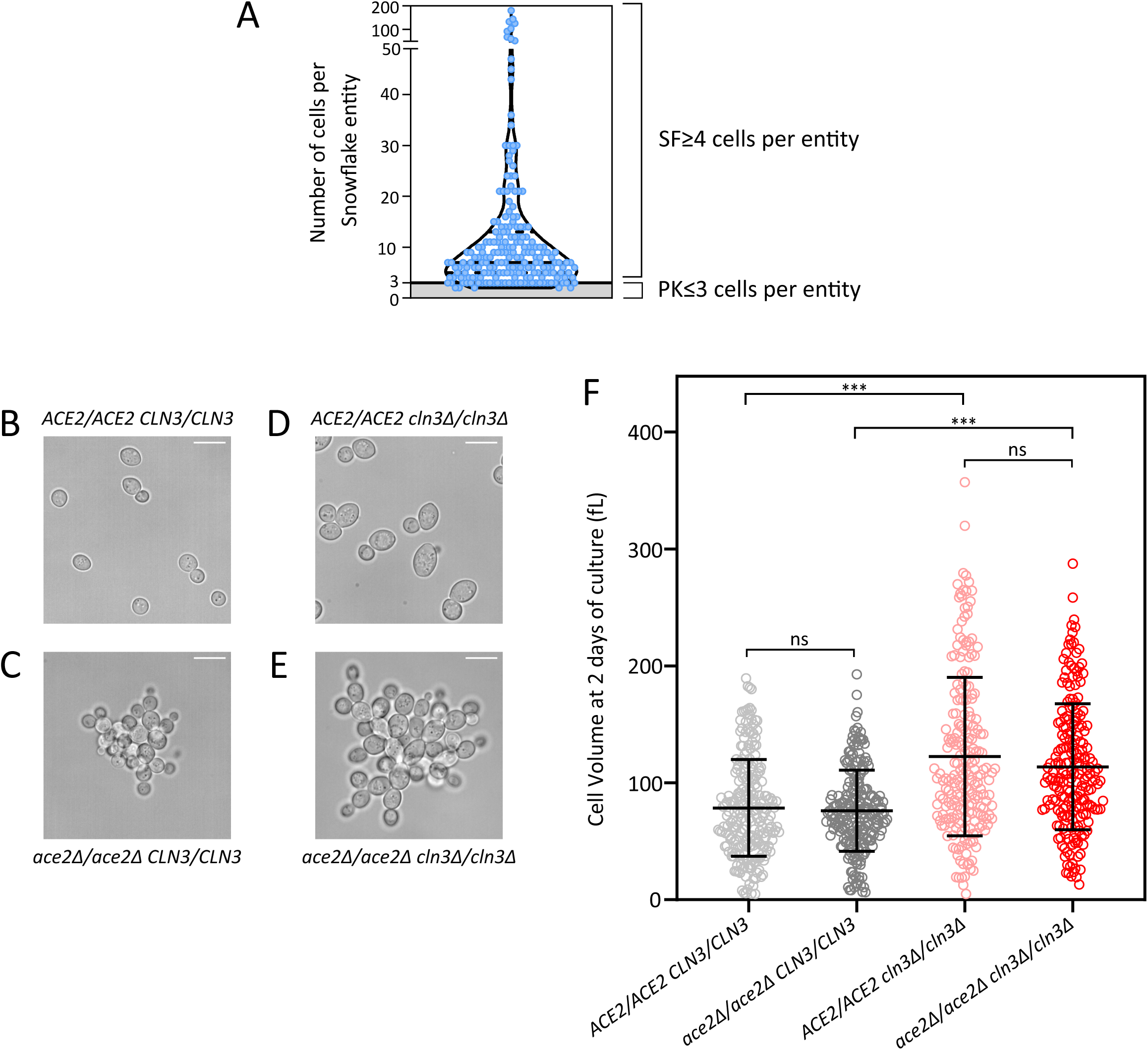
**(A)** Number of cells per Snowflake entity determined by counting the number of *HTB1*-*CFP* positive nuclei per Snowflake entity. **(B-E)** Representative images of *CLN3/CLN3* strains, either *ACE2/ACE2* **(B)** or *ace2/ace2* **(C)**, and *cln3/cln3* strains, either *ACE2/ACE2* **(D)** or *ace2/ace2* **(E)**. Bars are 10 µm. **(F)** Cell volume measured at 2 days of culture (error bars ± SD, unpaired t-test, Welch correction, ***p<0.001, ns p>0.05).

**Figure 2-figure supplement 1.**
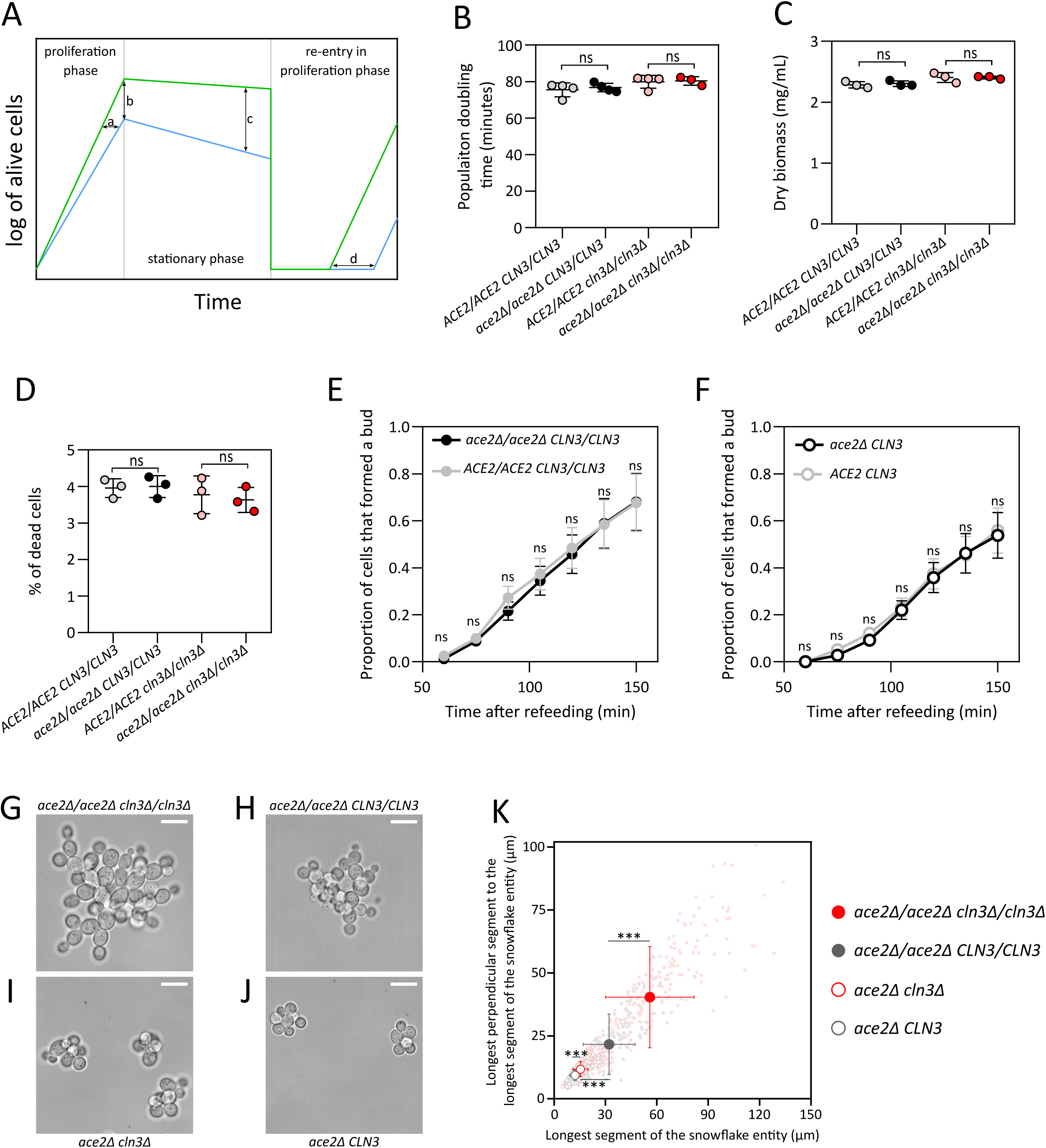
**(A)** Theoretical diagram showing the various phenomena that can explain, either alone or combined, how one subpopulation can outcompete another subpopulation in a co-culture. One subpopulation may divide more rapidly in exponential phase (a), may have a higher yield (b), may undergo less cell death during stationary phase (c) or may re-enter proliferation phase faster after refeeding (d). **(B)** Doubling time of populations in proliferation phase. Independent cultures (N>3) were maintained in constant proliferation phase for more than 24 hours, then the optical density at 600 nm was measured every 15 minutes for 8 hours (mean ± SD, unpaired t-test, Welch correction, ns p>0.05). **(C-D)** The dry biomass determined by filtration (**C**, N=3, mean ± SD, unpaired t-test, Welch correction, ns p>0.05) and the percentage of dead cells stained with ethylene blue (**D**, N=3, n>280, mean ± SD, Fisher’s exact test, ns p>0.05,) were also measured for 2 days stationary phase cultures. **(E)** Proportion of *CLN3/CLN3* cells that have formed a new bud after refeeding on a SDcasaWAU medium containing microscope pad of either *ace2/ace2* or *ACE2/ACE2* 2 days cultures (N=3, n>101, Fisher’s exact test, mean ± SD, ns p>0.05). **(F)** Proportion of *CLN3* cells that have formed a new bud after refeeding on a SDcasaWAU medium containing microscope pad of either *ace2* or *ACE2* 2 days cultures (N=3, n>91, Fisher’s exact test, mean ± SD, ns p>0.05). **(G-J)** Representative images of diploid *ace2/a*c*e2* snowflakes either *cln3/cln3* (**G**) or *CLN3/CLN3* (**H**), and haploid *ace2* snowflakes either *cln3* (**I**) or *CLN3* (**J**). Bars are 10 µm. **(K).** Size distribution of snowflakes in haploid (empty circles) or diploids (plain circles) backgrounds, either *CLN3* (black) or *cln3* (red). Size is estimated by measuring both the longest segment of the snowflake entity and the longest segment perpendicular to the first one (N=3, n>72). We compared the length of each segment independently with an unpaired t-test with a Welch correction (error bars ± SD, unpaired t-test, Welch correction, ***p<0.001, account for both segment).

**Figure 3-figure supplement 1.**
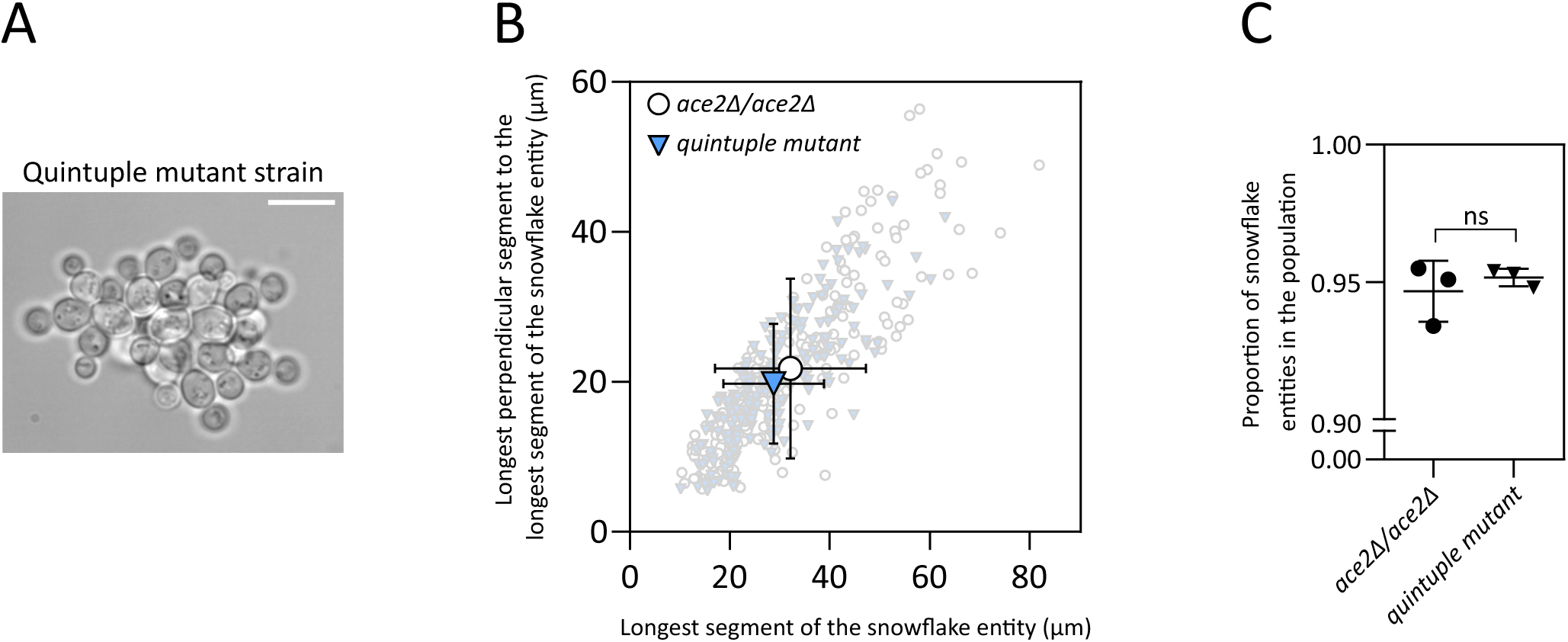
Snowflake characterization of quintuple mutant strains. **(A)** Representative image of the quintuple mutant strain. Bar is 10µm. **(B)** Size distribution of quintuple mutant (triangles) or *ace2/ace2* (circles) snowflakes. Size is estimated by measuring both the longest segment of the snowflake entity and the longest segment perpendicular to the first one. We compared the length of each segment independently with an unpaired t-test with a Welch correction (N=3, n>60, mean ± SD). **(C)** Proportion of snowflake entities in a population of quintuple mutant or *ace2/ace2* strains (N=3, n>201, mean ± SD, Fisher’s exact test, ns p>0.05).

**Figure 4-figure supplement 1.**
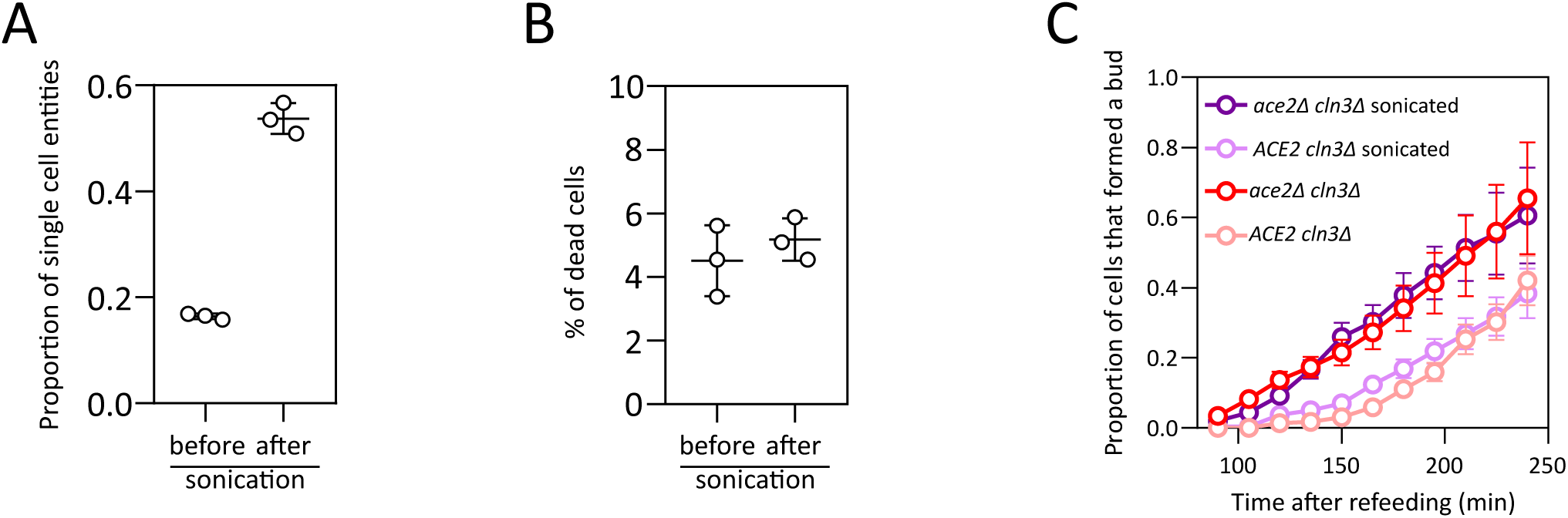
Effect of sonication on SF phenotype, cell death and quiescence exit. **(A-B)** The effect of sonication bath on an *ace2* culture was assessed by measuring the proportion of single cell entities (**A**, N=3, n>212) and the percentage of dead cells (**B**, N=3, n>207) in the population before and after the culture was sonicated. **(C)** Proportion of *cln3* cells, either *ace2* or *ACE2*, and either sonicated or not, that formed a new bud after refeeding of 2 days cultures on a SDcasaWAU medium containing microscope pad (N=3, n>63, Fisher’s exact test, mean ± SD).

**Figure 6-figure supplement 1.**
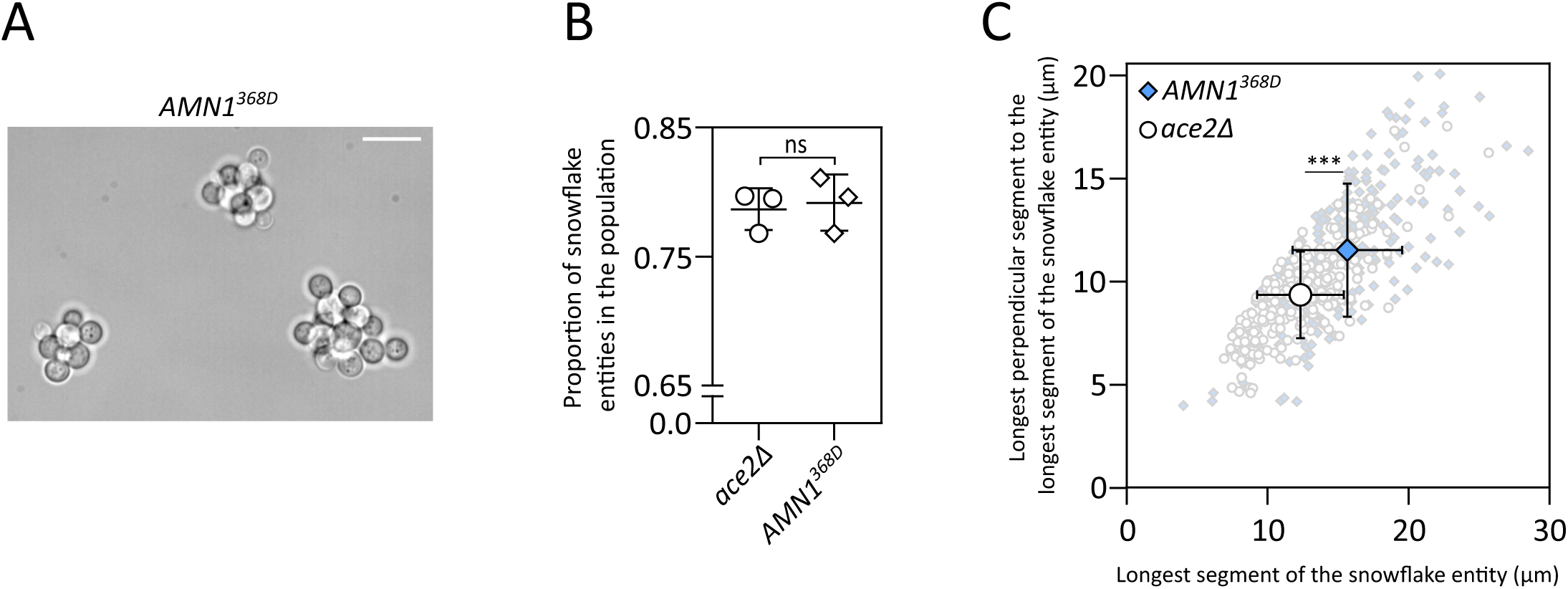
Snowflake characterization of *AMN1^368D^* strains. **(A)** Representative image of the *AMN1^368D^* strain. Bar is 10µm. **(B)** Proportion of snowflake entities in populations of *AMN1^368D^* or *ace2* haploid strains (N=3, n>219, mean ± SD, Fisher’s exact test, ns p>0.05). **(C)** Size distribution of *AMN1^368D^* (diamond) or *ace2* (circle) snowflakes. Size is estimated by measuring both the longest segment of the snowflake entity and the longest segment perpendicular to the first one (N=3, n>74, mean ± SD, unpaired t-test, Welch correction, ***p<0.001, account for both segment).

**Table S1:**
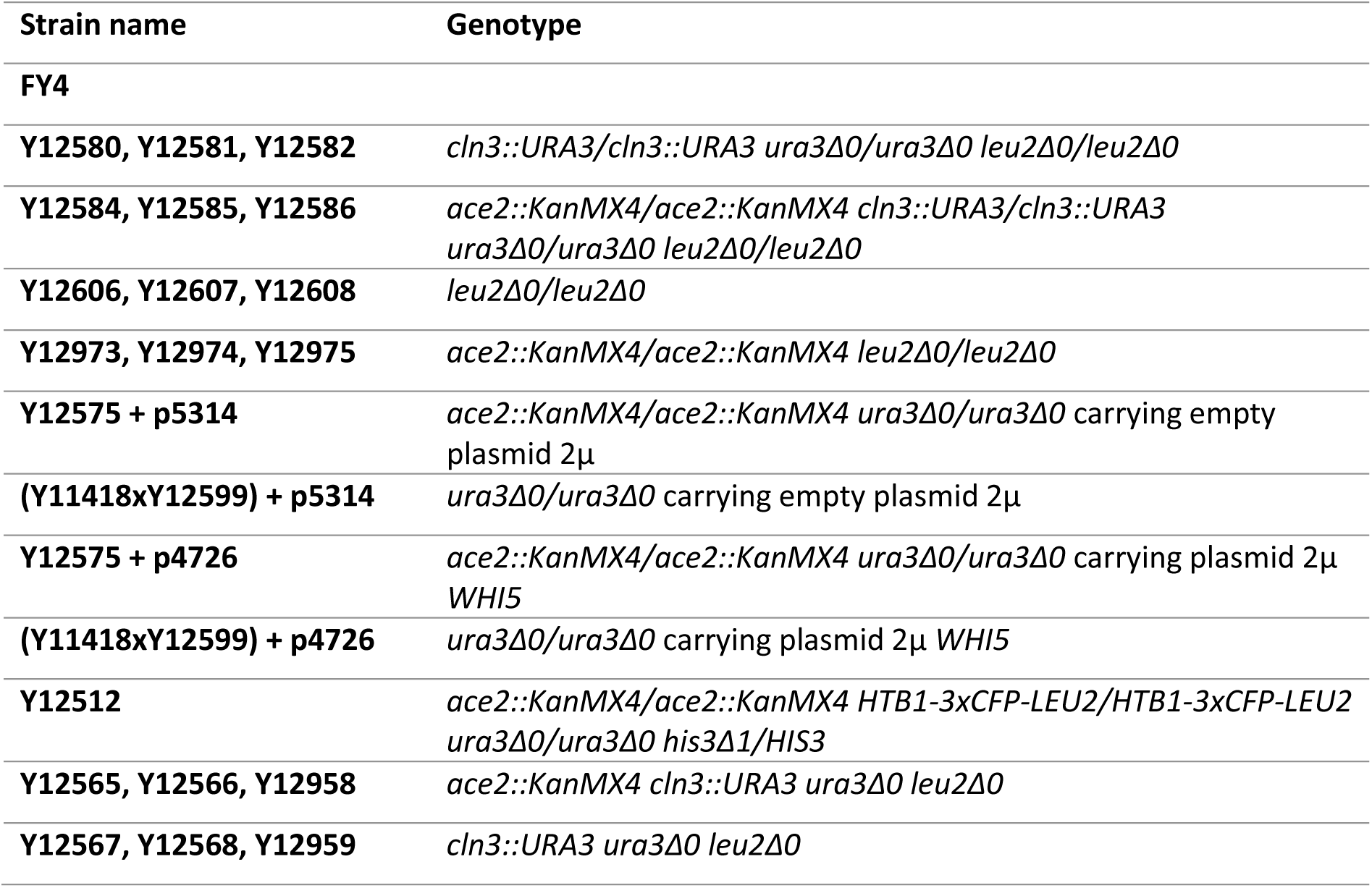
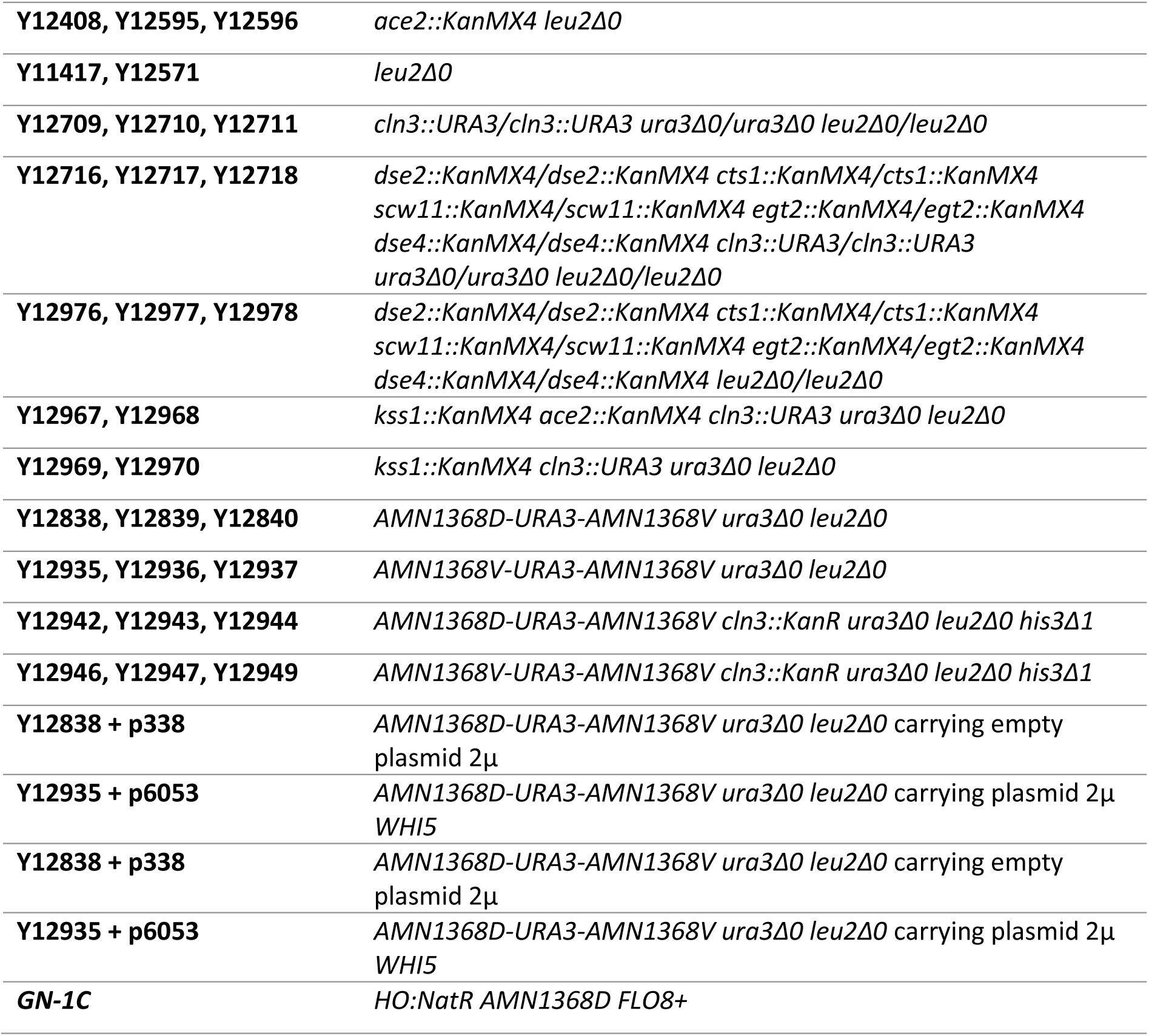
Strains.

**Table S2:** Plasmids.

| Plasmid Name | Description |  |
| --- | --- | --- |
| **p5314** | YEplac195: 2μ *URA3* AmpR | (35) |
| **p4726** | *WHI5* in YEplac195 | lab collection |
| **p338** | YEplac181 : 2μ *LEU2* AmpR | (35) |
| **p6053** | *WHI5* in YEplac181 | This study |
| **p6164** | YlpLac211 : integrative plasmid *URA3* AmpR | (35) |
| **p6165** | *AMN1-368D* in YlpLac211 | This study |
| **p6167** | *AMN1-368V* in YlpLac211 | This study |

**Table S3:**
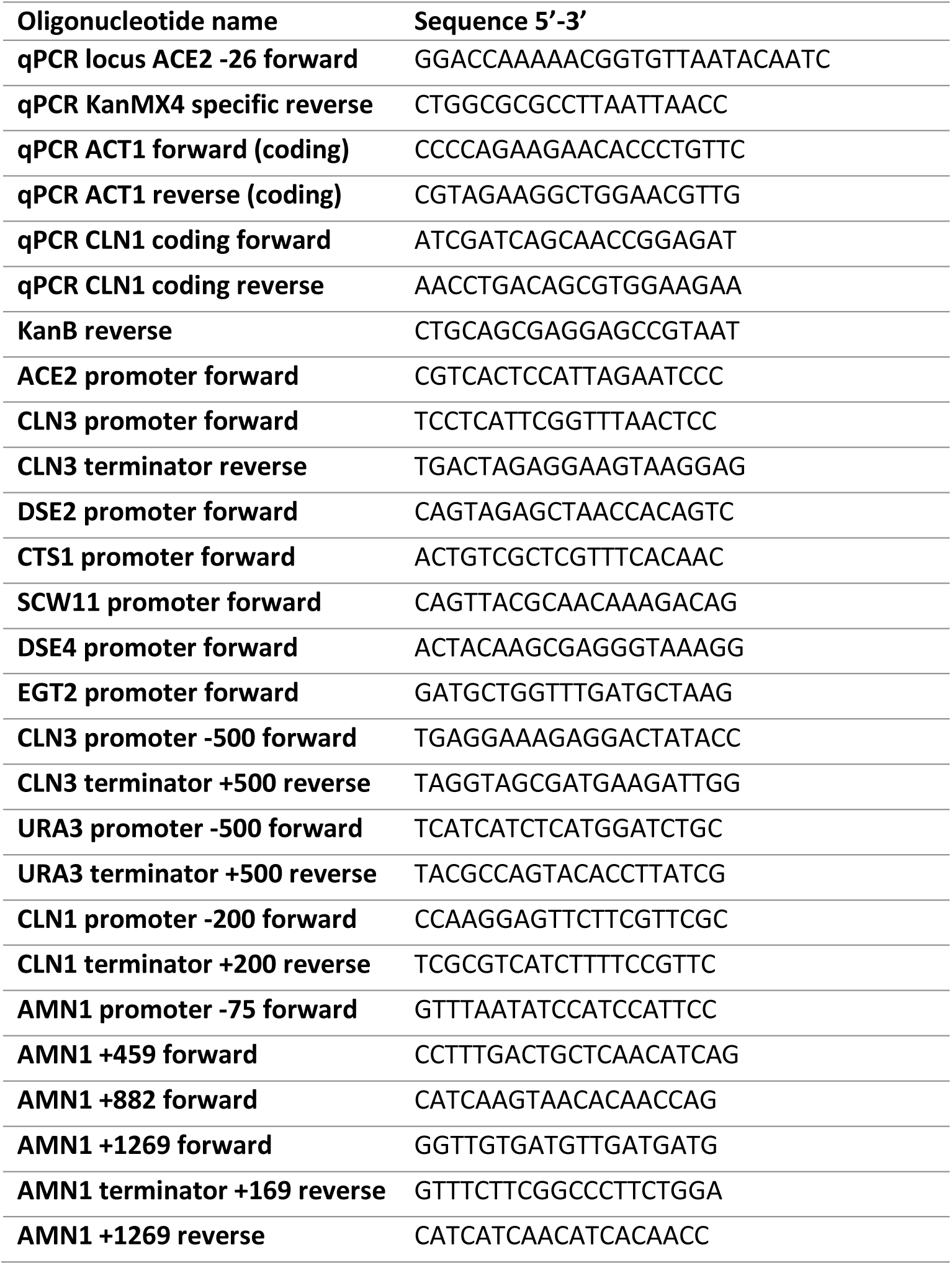
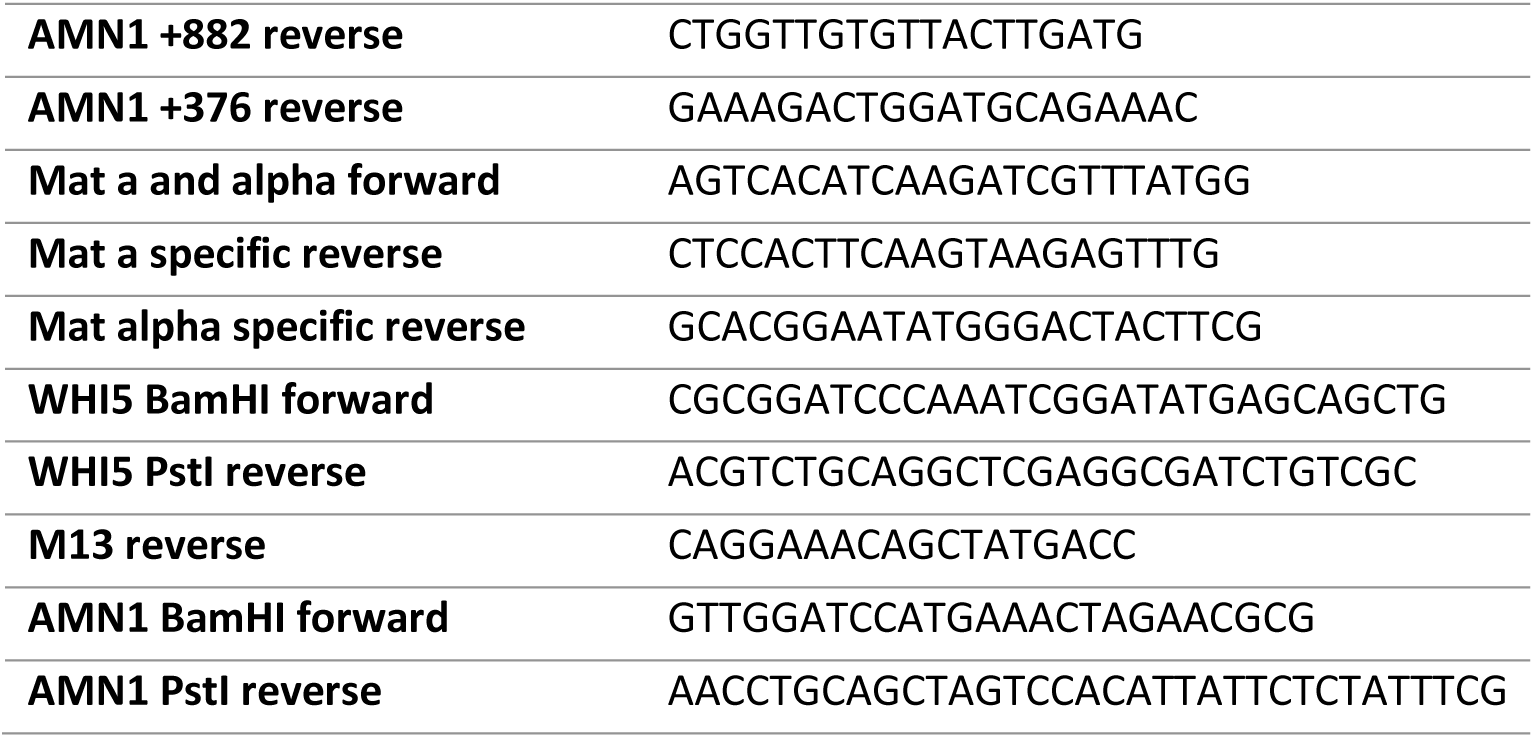
Oligonucleotides.

